# Modeling lipid nanoparticle transport in extracellular matrix: Effects of particle size and rigidity

**DOI:** 10.64898/2026.01.13.699279

**Authors:** Prasheel Nakate, Arezoo M. Ardekani

## Abstract

Lipid nanoparticles (LNPs) traverse through multiple biological barriers, such as crosslinked mesh structures in the extracellular matrix, before reaching their target sites. The physicochemical properties of LNPs determine their ability to penetrate complex biological environments such as the brain extracellular matrix. Their deformation in polymeric matrices affects transport, making it crucial to understand these factors for effective therapeutic delivery. Here, we develop a highly Coarse-Grained (CG) model of the LNP and its surrounding polymeric matrix, simulated as a uniform grid of cross-linked hyaluronic acid (HA) chains. The model for highly coarse-grained LNP was developed from a one-particle-thick membrane model that maintains mechanical features of lipid membranes, such as fluidity, topological changes, and hydrodynamic effects. Here, we investigate the collective influence of lipid nanoparticle size and its bending rigidity on the diffusive transport through the biological matrix. Our work highlights the role of particle to matrix size ratio in understanding deformation-assisted diffusive transport of LNPs in the matrix environment. Our study provides a tool to disentangle the effects of particle size and their bending rigidity on the transport through complex environments. Furthermore, this study systematically complements the rational design of lipid nanoparticle-based drug delivery platforms.

**SIGNIFICANCE:** Lipid nanoparticles (LNPs) are emerging as a powerful platform for delivering nucleic acid-based therapeutics, especially to hard-to-reach tissues like the brain. Their ability to protect and transport charged molecules, such as mRNA or siRNA, offers promising strategies for treating neurological disorders and advancing precision medicine for aging demographics around the world. In the brain parenchyma, LNPs must navigate the dense and heterogeneous extracellular matrix (ECM), composed primarily of crosslinked hyaluronic acid and proteoglycans. Understanding how particle size, deformability, and shape parameter affect the transport through this complex environment is critical for optimizing drug delivery platforms.

## INTRODUCTION

Lipid nanoparticles (LNPs) have played a crucial role in the delivery of mRNA drugs during the COVID-19 pandemic. In recent years, these nanoparticles have emerged as a promising tool for the delivery of a wide variety of therapeutic agents to the target sites. LNPs serve as a versatile nanocarrier platform for the effective transport and protection of hydrophobic and hydrophilic drugs such as proteins, nucleic acids, and small molecules in biological environments(1). One of the first LNP formulations, Doxil was developed for doxorubicin drug used for treating the ovarian cancer(2,3). Currently, the most extensive use of the lipid nanoparticle platform is found in cancer therapy. This is primarily associated with the increased solubility of hydrophobic drugs, improved drug circulation time, and reduced contact with the off-target sites as a result of the encapsulation(4,5). In addition to this, nucleic acid based therapeutics have shown great potential in treating various diseases(6). As, nucleic acids are highly negatively charged molecules they need to be encapsulated using the LNP platform for their delivery to the cells(7). Patisiran, also known as ONPATTRO, is one of the first approved siRNA drugs based on lipid nanoparticle platform(8, 9). Recently, the most notable successful application of lipid nanoparticles has been as delivery vehicles for the two newly approved COVID-19 mRNA vaccines(10). These vaccines deliver mRNA to cells, and their action eventually generates an immune response to the virus.

One of the main causes of mortality in aging demographics around the world is neurological disorders. Demographic analysts predict that by 2050, the global population of individuals over 60 years of age will reach 2 billion, with a significant portion exceeding 85 years of age(11). Despite a great understanding of genetic basis and pathology, age-related diseases such as Alzheimer’s and Parkinson’s disease have a low success rate in terms of drug development. Therefore, there is an urgent demand for the development of an effective platform for the delivery of therapeutic agents to the target sites in the brain(12, 13). Nucleic acid-based genetic medicines have shown great potential in the treatment of these neurological disorders. The site of action of these nucleic acids depends on their type. DNA-based medicines are generally delivered to the nuclei, whereas RNA-based therpeutics target the cytosol of the cell(14). Due to the presence of large negative charge on the nucleic acids they are prone to rapid clearance and poor biodistribution. Therefore, they are encapsulated using lipid nanoparticles that protect these therapeutic molecules before they reach target cells in brain tissue(15).

The extracellular space in brain tissue consists of a network of polymers known as the brain extracellular matrix (ECM). This matrix contains a crosslinked hyaluronic acid (HA) chain backbone along with proteoglycans, and tenascins(16). This structure is dynamic in nature as it can form a condensed network, freely float around cells, or be attached to the cellular surface(17,18). During brain drug delivery, lipid nanoparticles (LNPs) must overcome multiple biological barriers before reaching the brain parenchyma. Once there, they must navigate the dense extracellular matrix to reach the target cell membrane for effective delivery. This diffusive transport of nanoparticles is highly influenced by the physicochemical properties such as size, deformability, shape, and surface charge of nanoparticles (19, 20). A real-time multi-particle tracking (MPT) study conducted by Nance et al. showed that nanoparticles with a size as large as 114 nm were able to penetrate the human and rat brain tissues. Due to the dense PEG coating, these nanoparticles successfully navigated the extracellular barriers by evading the adhesive interactions with the matrix(21). Furthermore, this study showed that 72% of the pores in the extracellular space of human brain tissue are below 100 nm(21). In addition to size, the deformability of the nanoparticles also plays a significant role during their navigation through the extracellular barriers. In 2018, Yu et al. investigated the performance of rigidity-tuned core-shell nanoparticles through multiple intestinal and tumor barriers. Their findings indicated superior diffusivity of semi-elastic nanoparticles as compared to soft and hard core-shell nanoparticles(22). In another study, Yu et al. has shown that the temperature-induced changes in rigidity of liposomes affect their diffusive transport in the extracellular barriers. Here, the optimal diffusivity is observed at the phase transition temperature of the liposomes, indicating deformation-assisted rapid diffusion(23). In a recent review in 2024, Gadalla et al. surveyed the effects of nanoparticle deformability on transport across multiple biological processes, including circulation, extravasation, interactions with diverse cell types, movement through tissue matrices, and trafficking into intracellular organelles(24).

Despite advances in particle tracking techniques, accurately characterizing particle motion in vivo or ex vivo within complex environments such as the brain extracellular matrix remains a significant challenge. In particular, it remains extremely difficult to use particle tracking measurements to systematically investigate the effects of a wide array of physicochemical and mechanical properties of nanoparticles. In contrast, in silico modeling approaches, such as coarse-grained molecular dynamics simulations, provide a robust and flexible framework to study these properties across extensive parameter spaces. These simulations can provide time-dependent particle trajectories through complex microenvironments such as the brain extracellular matrix and effectively complement the experimental particle tracking techniques. In 2022, Hanssen and Malthe-Sørenssen employed a coarse-grained molecular dynamics study to examine single-particle diffusion through the perineuronal nets composed of negatively charged polymer brushes. This system consisted of simplified, unbranched polymer brushes, thereby motivating the need for a physiologically relevant, cross-linked polymeric structure for investigating nanoparticle transport in the brain extracellular matrix (25). Recently, He et al. utilized molecular dynamics simulations in conjunction with the single-particle tracking experiments to evaluate the effects of tumor ECM physical properties on the nanoparticle diffusion in the tumor microenvironment. These simulations enriched their findings by highlighting reduced nanoparticle diffusivity with increasing polymer network density and stiffness within tumors(26).

Transport processes of LNPs through the complex brain extracellular matrix encounter a multitude of physicochemical interactions, making it nearly impossible to determine the underlying diffusion mechanisms experimentally. Moreover, disentangling the effects of interdependent physicochemical properties within experimental systems presents an additional challenge. In this work, we have developed a coarse-grained molecular dynamics-based computational framework to investigate the impact of variable physicochemical properties of LNPs on the diffusive transport mechanism through the brain ECM. The matrix is modeled using a uniform crosslinked hyaluronic acid structure with Debye interactions for the screened electrostatic forces due to the presence of dissolved ions. Whereas, the LNP is represented by a one-particle thick, solvent-free membrane model developed by Yuan et al. (27). With this computational framework, we analyze the combined effect of LNP size relative to the matrix mesh size and the particle bending rigidity or deformability on the diffusive transport of neutral LNPs in the ECM. This study aims to provide critical insights into the role of physical properties on the particle transport and unravel the underlying diffusion mechanisms to better guide the design and optimization of lipid nanoparticle-based drug delivery platforms.

## METHODS

### Extracellular matrix model

To investigate the ability of lipid nanoparticles to diffuse through the extracellular matrix in the brain parenchyma, we construct a cross-linked structure representing hyaluronic acid (HA) chains. Here, we simplify the matrix structure by keeping four crosslinking sites in each direction at uniform distances. This structure consists of 48 HA chains, with each chain represented by 49 beads. The structure has 64 cross-linking site beads that hold the grid anchored in all directions. Here, we consider a periodic simulation box of size 98*σ* × 98*σ* × 98*σ* for most of the cases (*σ* is the unit length) with the distance between adjacent crosslinking sites as 24*σ*. The units of mass, time, and energy are *m, τ*, and *ε*, respectively. HA molecules consist of negatively charged repeating disaccharide units that describe the unit mass *m* in the system. We represent each disaccharide unit by one coarse-grained bead with a charge of *-e*. These beads are connected with each other through covalent bonds modeled as harmonic stretching potentials. Along with this, we add harmonic bond potentials between the adjacent bonds to maintain the structural integrity of the chain. In this study, we add restrained harmonic potentials on the beads at the crosslinking sites as we focus on the motion of lipid nanoparticles through this matrix structure(22,23). These restraining potentials tether the crosslinking sites to their initial positions, which represent the stationary extracellular matrix interacting with the lipid nanoparticles(26). To parameterize the potentials for chains in this model, we refer to the study done by Yu et al. on a similar physiological matrix structure that interacts with nanoparticles(22). Following this study, we assign the equilibrium bond length for these beads as 2*σ*, and the stiffness constant is kept at 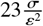 (22). The cross-linked nodal beads are tethered to their initial positions with a stiffness constant of 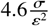 (22). The bond angles of two adjacent bonds are constrained by harmonic bond potentials with an equilibrium bond angle of 180 ^°^ and a stiffness constant of 4.6*ε* (22).

The non-bonded interaction between the HA beads is modeled using the Lennard-Jones potential shown in equation 1

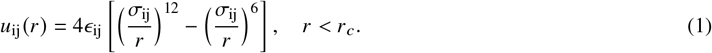

Here, *ϵ* is the strength of interaction between the beads and *r* is the distance between them. *σ*_*ij*_ is the equilibrium separation distance between two beads and *r*_*c*_ is the distance at which the potential is cutoff. The matrix structure in the present study is stationary in nature. Due to the restrained potentials on crosslinking sites, the chains never undergo translational motion in the domain. Therefore, the probability of active non-bonded interactions between the chain beads is significantly reduced. As a result, the interaction strength *ϵ* between the chained HA beads is not influential in this model. Similar to the bonded interactions from the study of Yu et al., we assign the non-bonded interaction between the chained HA beads as *ϵ*_*ij*_ = 0.1(22). The lipid nanoparticles in this study are non-aggregating in nature; therefore, we model the interaction strength as *ϵ*_*ij*_ = 0.01, which corresponds to the least attractive interactions in the model. To assign the interaction strength between the LNP bead and the chained HA beads, we perform a parametric study shown in Figure 4. Here, we test the model performance for *ϵ* values ranging from 0.02 to 0.05. All of these values are listed in Table 1.

**Table 1:** Lennard-Jones interaction parameters for the LNP beads and the hyaluronic acid beads.

| Bead type i | Bead type j | $\epsilon_{ij}(\epsilon)$ | $\sigma_{ij}(\sigma)$ | $r_c(\sigma)$ |
| --- | --- | --- | --- | --- |
| LNP | LNP | 0.01 | 1.0 | 2.0 |
| HA | HA | 0.1 | 1.0 | 2.0 |
| LNP | HA | 0.02 - 0.05 | 2.0 | 5.0 |

Additionally, these beads have long-range electrostatic interactions because of the negative charges present on the disaccharide units of the HA. These negative charges are screened by the dissolved ions present in the extracellular matrix. Therefore, to model electrostatic interactions, we utilize the Debye potential shown in equation 2

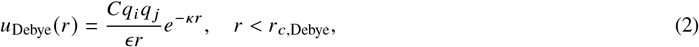

where *q* is the charge on the bead and *ϵ* is the dielectric permittivity. *κ* is the inverse of the Debye length, *C* is the energy conversion constant, and *r*_*c*,*Debye*_ is the cutoff for this Debye potential. The brain extracellular fluid has a greater proportion of the monovalent ions, such as sodium *N a*^+^ and chlorine *Cl*^−^, as compared to the divalent ions(28). Based on the ionic strength of typical interstitial fluid in the extracellular space, the Debye length for monovalent ions is ~ 1 nm(29). Recently, Hanssen and Malthe-Sørenssen also used a similar value of Debye length in their study involving perineuronal net structures(25).

### Model for Lipid nanoparticles

To model full-size lipid nanoparticles, we utilize a one-particle-thick, solvent-free membrane model developed by Yuan et al.(27). Such a highly coarse representation of the lipid molecule pushes the limits of the spatiotemporal scales of the computational problem to the maximum extent. This model successfully captures the essential mechanical features of lipid membranes, such as fluidity, topological changes, and hydrodynamic effects (27). Due to a highly coarse representation of the lipid beads, this model features a soft-core interaction between the particles as compared to the Lennard-Jones potential. Here, we model the interparticle interaction between lipid beads using a combination of distance-dependent *u* (*r*) and orientation-dependent 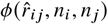 functions as described by Yuan *et al*.(27).

The distance-dependent interparticle potential *u* (*r*) has a repulsive *u*_*R*_ (*r*) and an attractive branch *u*_*A*_ (*r*) that meet at *r* = *r*_*min*_. Here, *r*_*min*_ = 2^1/6^*σ* is the distance at which the potential reaches a minimum. The repulsive branch shown in equation 3 has the form of a 4-2 LJ potential, unlike much steeper 12-6 LJ potential. Whereas the attractive branch in the equation 4 has a cosine function that approaches zero at the cutoff distance *r*_*c*_. The exponent *ξ* is used to tune the slope of this branch, which subsequently modulates the in-plane diffusivity of particles. The distance-dependent potential in the model is described as follows

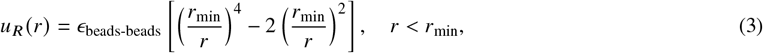

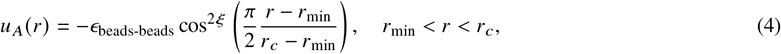

The orientation-dependent functions 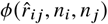 and 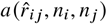 are used to model the hydrophobic effect, and they are described as

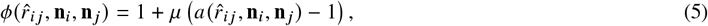

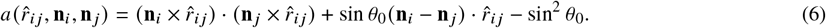

Here, **n**_*i*_ and **n**_*j*_ are the unit vectors that represent the symmetry axes of the bead *i* and *j*, respectively, and 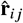 is the unit vector pointing from bead *j* to *i*. Finally, the distance-dependent and orientation-dependent functions are combined, which leads to a complete form of anisotropic potential as shown below

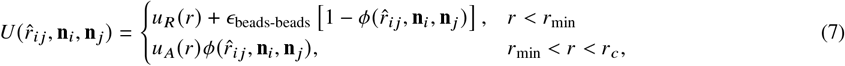

In this form, the function 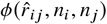 from equation 5 adds an additional term to shift the repulsive branch *u*_*R*_ (*r*) and scales the attractive branch *u*_*A*_ (*r*). This treatment ensures variable interaction strengths for different relative orientations. The parameters *θ*_0_ from the equation 6 and *μ* from the equation 5 correlate with the spontaneous curvature and the bending rigidity of the lipid membrane, respectively. The spontaneous curvature *c*_0_ of the membrane is inversely proportional to the radius of the preassembled spherical lipid nanoparticle *r*_0_, and it is related to *θ*_0_ by the expression, 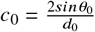, where *d*_0_ is the mean interparticle distance (27). Based on this relationship, we choose 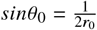, with mean interparticle distance of 1*σ*. As the size of LNP increases, the sin *θ*_0_ parameter decreases. In this study, we chose the same parameters for the lipid membrane as reference (27), *ϵ*_beads-beads_ = 1*ε, ξ* = 4, and *r*_*c*_ = 2.6*σ*.

The model parameter *μ* in this system controls the orientation-dependent function 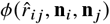 in equation 5. When particle orientation reaches its most favorable value, i.e, *θ*_*i*_ = *θ*_*j*_ = *θ*_0_, then the values of 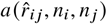 and 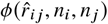 become 1, and the anisotropic potential in the equation 7 is left with only distance-dependent terms. But when this particle orientation deviates from *θ*_0_, the system tries to bring it back to its energetically most favorable shape. Here, the model parameter *μ* decides the degree of energy penalty given to these particles. As this penalty increases, it gets harder for particles to change their favorable orientation *θ*_0_, which subsequently increases the bending rigidity of the LNP structure. In 2010, Yuan et al. reported a monotonically increasing parametric relationship between the membrane bending rigidity and the parameter *μ*, as shown in Figure S2 of the supplementary material. The reported bending rigidity values range from 12*k*_*B*_*T* to 40*k*_*B*_*T*, that aligns with the experimental data. Recently, Chakraborty et al. estimated the bending rigidity of the lipid vesicles with variable cholesterol content using neutron spin-echo (NSE) spectroscopy. In these experiments, the bending rigidity values ranged from 19*k*_*B*_*T* for 0% cholesterol to 57*k*_*B*_*T* for 50% cholesterol (30). Based on these studies and the parametric curve in Figure S2 of the supplementary material, we choose three representative values of bending rigidities of different lipid nanoparticle types soft, semi-elastic, and hard, as summarized in Table 2.

**Table 2:** Model parameters for LNPs having different bending rigidities for sizes 12*σ*, 16*σ*, and 20*σ*.

| | $\mu$ | Bending rigidity ($k_B T$) |
| --- | --- | --- |
| Soft LNPs | 3 | 23 |
| Semi-elastic LNPs | 5 | 37 |
| Hard LNPs | 8 | 45 |

### Simulation details

We perform coarse-grained molecular dynamics simulations using the LAMMPS simulation package. We introduce 10 pre-assembled lipid nanoparticles in the extracellular matrix model to study the diffusive transport of these LNPs through the matrix. The beads of the LNPs interact with the chained beads of the HA matrix through the Lennard-Jones potential as shown in equation 1. The interaction parameters for the potential between these bead types are listed in Table 1. The equations of motion are integrated with the Velocity-Verlet algorithm. Here, we keep the timestep *δt* = 0.01*τ*, where 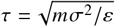 is the time unit.

Here, we employ the Langevin thermostat along with the NVE microcanonical ensemble, where the total number of particles *N*, the system’s volume *V*, and the total energy *E* remain constant for both LNP and extracellular matrix beads. The system temperature is kept at *k*_*B*_*T* = 0.23*ε*, where *k*_*B*_ is the Boltzmann constant. For all the cases, the total length of the simulation is kept at 2 × 10^5^*τ*, and bead position data is stored at every 100*τ*. To test the effect of domain size on the LNP diffusive behavior in the matrix, we perform simulations for five sizes ranging from 98 to 194 *σ* with an increment of 24 *σ*. Here, the number of nodal crosslinking beads in each direction increases from 4 to 8. For all five cases, the number density of LNPs is kept constant. Additional details for these tests are provided in the supplementary material. To quantify the motion of particles in the matrix, we compute the mean square displacement (MSD) of the LNP centroid as it traverses the matrix structure using the equation 8

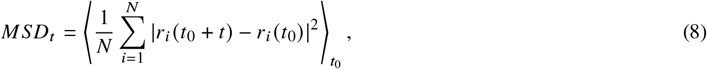

where *N* is the number of LNPs in the system, *r* represents the coordinates of the centroid of LNPs, *t* is the time lag, and *t*_0_ is the time origin. Figure 3 shows the MSD of the LNPs for increasing domain sizes. These trends indicate that domain size has a negligible effect on the overall motion of the LNPs. Here, the smallest box has 10 LNPs, whereas the largest box of size 194 *σ* has 80 LNPs. These plots suggest that the smallest domain of 98 *σ* can sufficiently capture the overall statistics of the system for diffusive transport analysis.

## RESULTS AND DISCUSSION

### Effect of particle-matrix relative affinity

The degree of affinity between the LNP beads and the HA matrix polymeric beads depends on the nature of the interaction between the chemical entities on the LNP surface and the matrix structure. In the coarse-grained modeling framework, the Lennard-Jones potential parameter *ϵ*_*i j*_ tunes the interaction between the LNP and the matrix structure. Typically, a higher *ϵ*_*i j*_ value corresponds to a deeper potential well and stronger attractive interactions, and vice versa. To test the model’s performance for variable affinities, we simulate the system with four different values of *ϵ*_*i j*_ ranging from 0.02 to 0.05. Figure 4 a) shows the mean square displacements of LNPs for different values of particle-matrix affinities. Along with this we plot the trajectory data of a single LNP through the matrix for *ϵ* = 0.02 and *ϵ* = 0.05 in Figures 4 b) and 4 c), respectively. These results suggest that as the affinity parameter increases, the motion of LNPs in the matrix is more restricted, and they tend to bind with the matrix structure. In this study, we choose *ϵ* = 0.02 to represent LNPs with the least attractive interaction with the matrix structure, allowing us to isolate and examine the effects of particle size and rigidity on their diffusive behavior in the matrix.

### Effect of particle size relative to mesh size

The pore size distribution obtained by Nance et al. suggests that 72% of the brain extracellular space consists of pores smaller than 100 nm(21). This limitation presents a significant challenge in the delivery of larger LNPs to the target sites in the brain. Reduction in the size of LNPs results in lower drug volume inside the LNP core. Additionally, there are significant process-dependent challenges in manufacturing small size LNPs at scale. To analyze the impact of LNP size on their transport in the extracellular space, we define size ratio (SR) as a ratio of LNP size to the crosslinked mesh size in the hyaluronic acid matrix structure shown in Figure 1. Here, we perform simulations for size ratios 50%, 67%, and 84% corresponding to the LNP sizes 12*σ*, 16*σ*, 20*σ* with a constant crosslinked mesh size of 24*σ*. Based on the length scale of the current model, this crosslinked mesh size approximately corresponds to 72-96 nm. This estimate falls below the 100 nm threshold pore size described in the study of Nance et al.(21). Therefore, based on the size ratios considered in this study, the lipid nanoparticle model dimensions fall in the range of 48-80 nm. This range matches well with the nanoparticle size ratios used in the study performed by McKenna et al. in the realistic biological environment of rat brain slices(31). Figure 5 a) shows the snapshots of the simulations with varying size ratios of LNPs. Figure 5 b) displays the trajectory of a single LNP traversing in the hyaluronic acid matrix for each size ratio. These plots indicate that the trajectory of smaller LNPs is more spread out in the matrix, whereas the LNPs with larger size ratios experience greater hindrance from the matrix structure. The motion of LNPs in the matrix alternates between the confinement and breakout regions. These areas can be easily distinguished in the case with a 50% size ratio, where confined and breakout regions are highlighted with black and blue colors, respectively. As the particle size ratio increases, the breakout regions in the matrix reduce, and the confinement region grows larger, evident in the cases with 67% and 84% size ratios.

**Figure 1.**
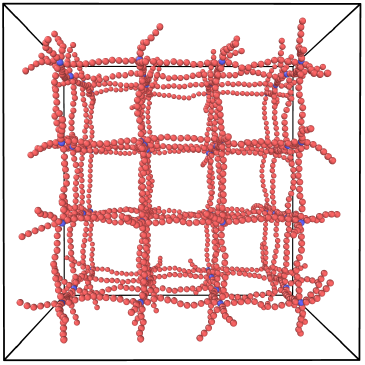
Uniform grid network of hyaluronic acid polymer chains

**Figure 2.**
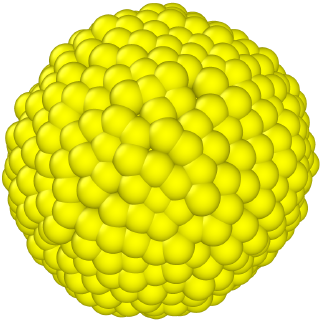
Lipid nanoparticle assembled with the coarse-grained beads using one-particle thick membrane model.

**Figure 3.**
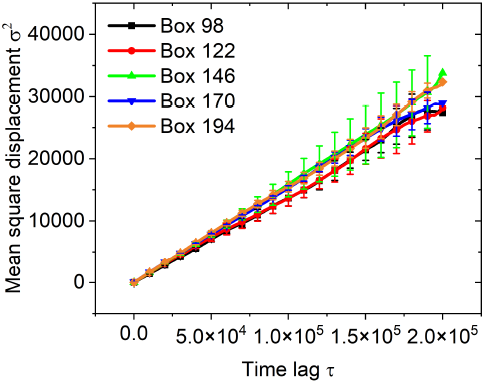
Mean square displacements of LNPs in the extracellular matrix with increasing domain size.

**Figure 4.**
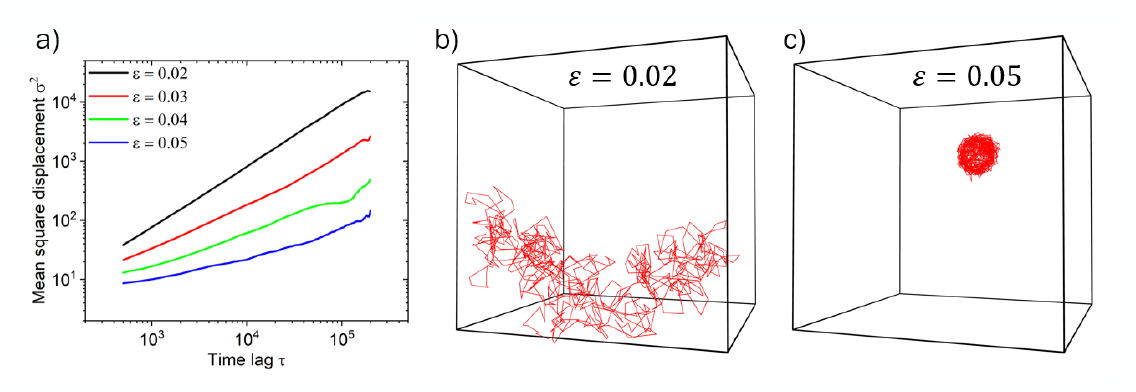
Effect of variable particle-matrix relative affinity on a) Mean square displacements, and b), c) trajectory of the LNPs through the extracellular matrix.

**Figure 5.**
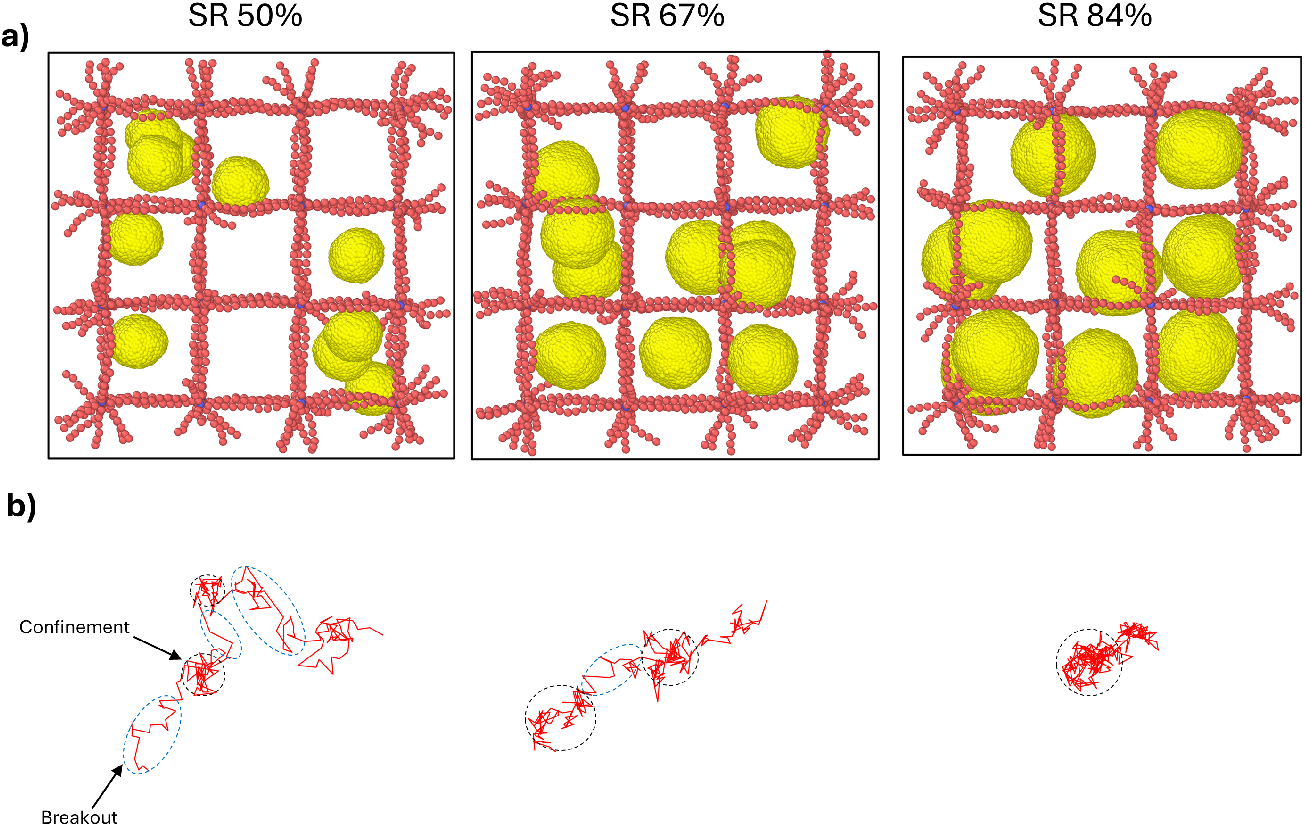
a) Simulation snapshots and b) trajectories of LNPs with areas highlighting confinement and breakout regions for increasing LNP to matrix mesh size ratio

For quantitative comparison, we calculate MSDs of semi-elastic particles having different size ratios in Figure 6 a). All of these MSDs and their standard deviations are obtained from atleast three independent simulation runs with random initial configuration of LNPs in the matrix. These trends suggest that particles with 50% size ratio effectively diffuse in the matrix structure as compared to 67% and 84% size ratios. As the size ratio increases, the degree of restriction from the matrix increases in a nonlinear manner. All three MSD curves indicate that the particles undergo normal diffusion in the matrix, where MSD increases almost linearly with the time lag. To further quantify the performance of LNPs in the hyluronic acid matrix, we evaluate the diffusivity of LNPs. We obtain these values by fitting the portion of the MSD roughly between 50,000 *τ* to 150,000 *τ*. This approach avoids the anomalous behavior of LNPs at shorter timescales and the higher variations due to small sample size at longer timescales. Additional details for fitting the MSD curve are provided in the supplementary material. Figure 6 b) shows the diffusivity of the LNPs in scaled units (*σ*^2^/*τ*) extracted from the slope of the linear fit in Figure 6 a). The diffusivity of LNPs at a size ratio of 84% is reduced by nearly threefold compared to the 67% case, and by sevenfold relative to the 50% case. This trend indicates a non-linear nature of decline in the diffusivity as the size ratio increases.

**Figure 6.**
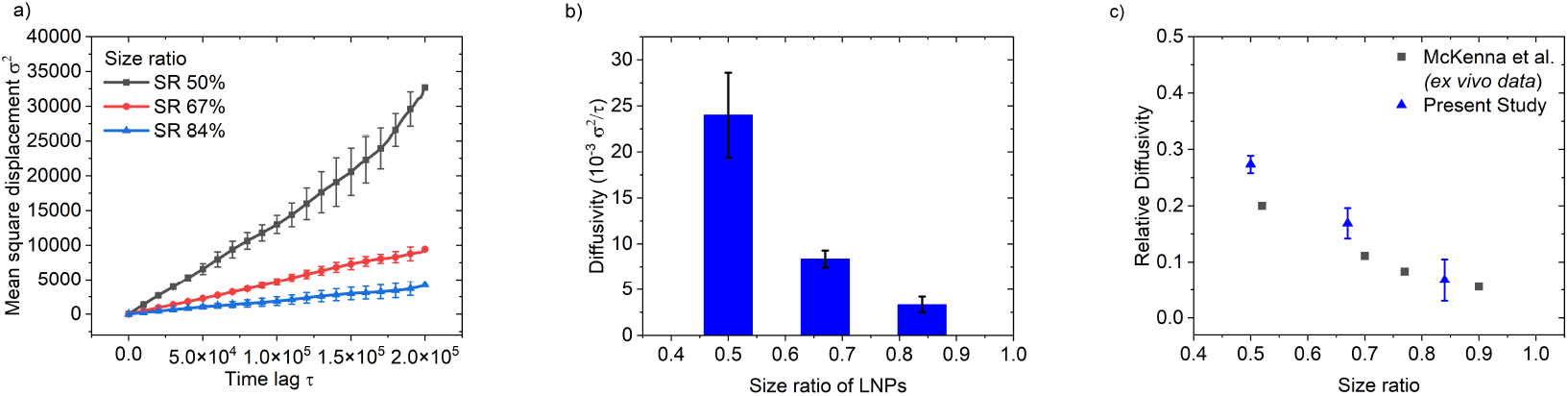
Effect of particle size relative to mesh size: a) Mean square displacements of semi-elastic LNPs, b) Diffusivity of semi-elastic LNPs for variable size ratios in the matrix, and c) Size ratio dependent relative diffusivity of hard LNPs in the matrix compared against *ex-vivo* multiparticle tracking data of McKenna et al. (31).

To test the model performance against experimental observations, we utilize a validation strategy based on relative diffusivity(32). This is calculated as a ratio of the diffusivity of nanoparticles in the matrix to the diffusivity in water. This is a standard metric commonly used to compare the results across different methods (33), as most of the quantities involved in modeling framework are unitless(23). Recently, He et al. developed a model for diffusion of nanoparticles in tumor extracellular matrix, where the model is validated using unitless quantity relative diffusivity from experimental data on silica nanoparticles(26). In this work, we validate the model performance against the experimental data on diffusivity of nanoparticles in the rat brain extracellular matrix(31). McKenna et al. performed multi-particle tracking analysis for 40 nm PS-PEG nanoparticles in different rat brain slices with average pore sizes ranging from 44 nm to 77 nm that covers the range of size ratios from 52% to 90%. As the PS nanoparticles are inherently rigid, we use relative diffusivity data from the hard LNPs to compare the model performance. Additional details for calculating relative diffusivity are mentioned in the supplementary materials. Figure 6 c) shows the trend in the relative diffusivities of LNPs with increasing size ratios. Our simulations with size ratios ranging from 50% to 84% compare qualitatively well against the relative diffusivity obtained from the *ex vivo* study of McKenna et al. We observe some deviations in Figure 6 c) towards lower size ratios as the multi-particle tracking data is obtained from a realistic biological environment where there is spatial heterogeneity in the mesh sizes that evolve dynamically. The comparison between the relative diffusivity of nanoparticles is limited by differences in the precise mechanical properties of the nanoparticles used in this study versus those in the experimental system, which are generally more rigid. Nonetheless, the qualitative agreement observed between the hard LNPs in this simulation and the nanoparticles in the study of McKenna et al. is promising.

### Effect of particle rigidity on the diffusivity through the matrix

Particle deformability has a great impact on the diffusive transport of LNPs through the HA matrix. The effect of deformability is highly correlated with the particle size ratios. As LNP surface area increases, the total number of deformation modes also increases. In order to understand this correlation, we perform simulations for particle rigidities 23 *k*_*B*_*T*, 37 *k*_*B*_*T*, and 45 *k*_*B*_*T* for each size ratio as shown in the Table 2. The bending rigidity of LNPs is tuned using the model parameter *μ* that assigns a penalty to the deformation of the membrane surface of the LNP based on its magnitude(27).

#### Particle Size ratio: 50%

Figure 7 a) shows the mean square displacements for three particle types: soft, semi-elastic, and hard. We calculate these MSDs from atleast three independent simulation runs for each particle type to reduce the non-linearities in the MSD curve. The ensemble behavior of the mean square displacements of all three types evolve in the same way. For the shorter time lags below 3 × 10^4^, the trends are indistinguishable, and they approach to similar displacements at longer time lags. Here, all the particle types have almost the same diffusivity as shown in Figure 7 b). This suggests that the bending rigidity of LNPs for a size ratio of 50% has almost no effect on their diffusive performance through the matrix structure. Here, the volume occupied by the ten LNPs is approximately 0.96% of the total system volume. This indicates that there is a significantly large space for the LNPs to explore, and any effect related to particle deformability is insignificant.

**Figure 7.**
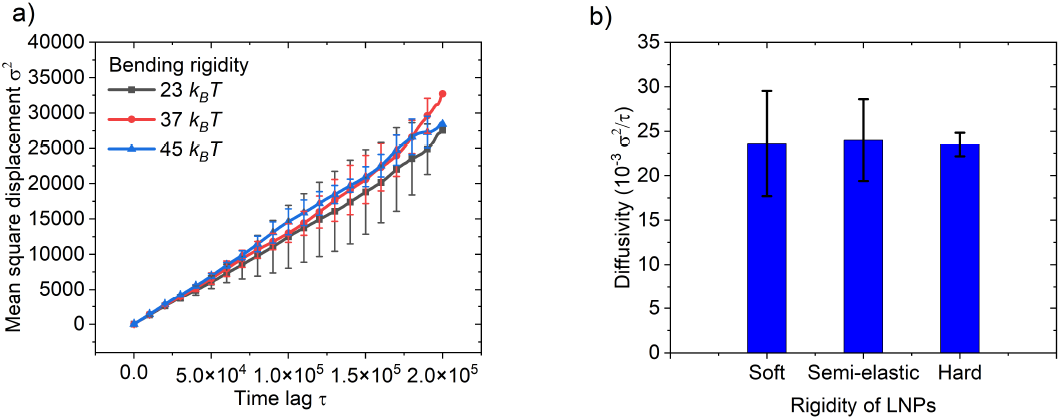
a) Mean square displacements and b) Diffusivity of LNPs with variable rigidities for a size ratio of 50%.

To further investigate this finding, we calculate the shape descriptor of LNPs using the gyration tensor. Here, we compute the asphericity parameter *b*, which measures the deviation of LNPs from the perfect spherically symmetric shape as follows,

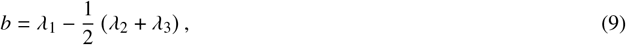

where *λ*_1_ ≥ *λ*_2_ ≥ *λ*_3_ are the three eigenvalues of gyration tensor(34). When all three eigenvalues are equal, the particle shape is perfectly symmetric. LNPs have a fluid membrane that dynamically fluctuates, resulting in an asymmetrical shape. Additional details for calculation of aphericity parameter *b* and their moving time averages are provided in the supplementary material. Figure 8 shows the evolution of the mean asphericity of LNPs and their moving time average during the simulation. Here, for all the LNP types with a 50% size ratio, asphericity values are above 1, indicating deviation from the perfectly spherical shape.

**Figure 8.**
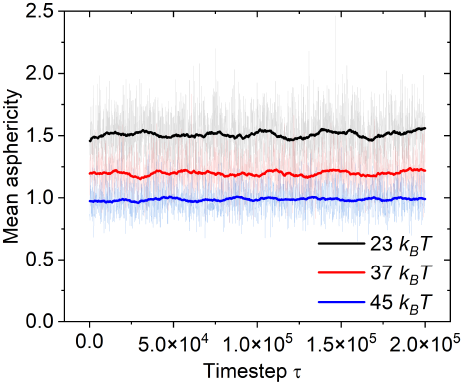
Mean asphericity of LNPs with different rigidities across the simulation time for particle size ratio 50%.

Notably, the mean asphericity of soft LNPs (23 *k*_*B*_*T*) is about 1.5, and that of hard LNPs (45 *k*_*B*_*T*) is 1. This indeed suggests that softer LNPs deform more readily in the matrix. However, the overall variation in mean asphericity among different LNP types is about 0.5, and it is insufficient to produce significant changes in the mean square displacement and diffusivity of LNPs within the matrix.

#### Particle Size ratio: 67%

To further test the effect of particle deformability on the diffusive transport through the matrix, we consider particles with a size ratio of 67%. Here, the LNPs occupy approximately 2.28% of the total system volume, with a size of 16*σ*. Figure 9 a) demonstrates the evolution of mean square displacements of LNPs with all three types of bending rigidities. Similar to the previously mentioned cases, we calculate the MSDs from atleast three independent runs that initiate from randomly assigned LNP configurations in the matrix. Here, all the three LNP types show almost similar MSDs as they traverse through the matrix. The MSD curves for all bending rigidiy cases evolve almost parallel to each other which reaches to final square displacement around 10,000 *σ*^2^. As a result, we observe the mean diffusivity values in the range of 7.23 × 10^−3^ − 8.36 × 10^−3^*σ*^2^/*τ* shown in the bar plot of Figure 9 b). This indicates that similar to 50% size ratio case, bending rigidity of LNPs have no effect on their diffusive performace at size ratio 67%.

**Figure 9.**
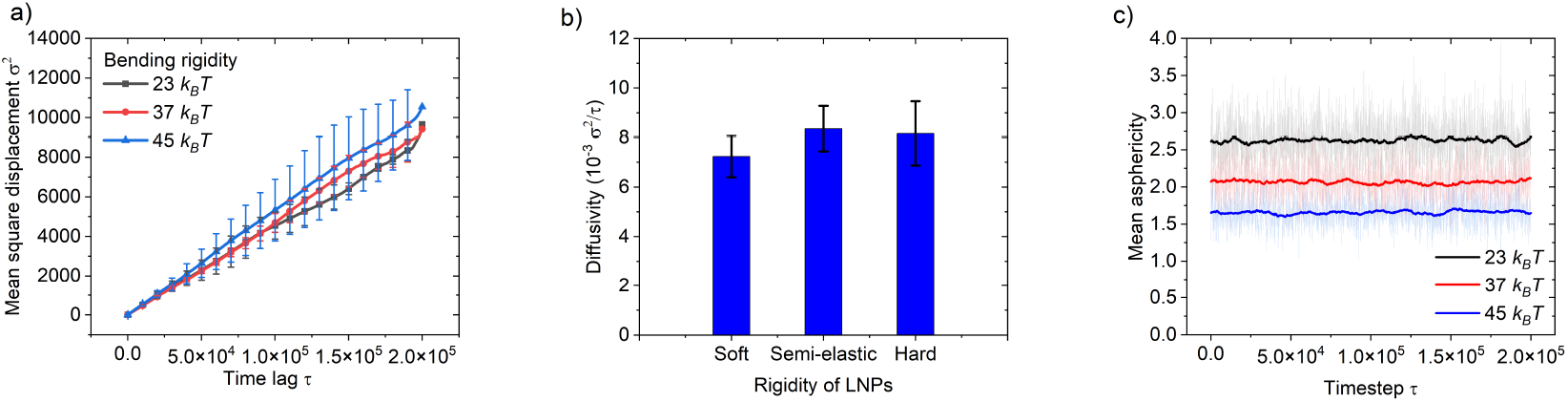
a) Mean square displacements, b) Diffusivity of LNPs with variable rigidities, and c) Mean asphericities of all LNP types for the size ratio of 67%.

To analyse LNP shape fluctuations, we plot the mean asphericities of all LNP types in the Figure 9 c). Here, all three asphericity values are much greater than 0, showing more deviation from spherical shape as compared to LNPs in 50% size ratio cases. The highest asphericity of LNP in this case is around 2.6, indicating significant deformation of the softer LNPs. The asphericity data presented in Figures 8 and 9 c) suggest that the LNP shape fluctuations within the 50% and 67% size ratio cases are not sufficient enough to measurably influence the trajectory and overall displacements of LNPs in the matrix environment. The LNP shape fluctuations can be increased further by reducing the membrane bending rigidity parameter *μ* or increasing the size of the LNPs. But there is a threshold value for rigidity parameter *μ* below which the orientational dependence in the model becomes weaker and membrane starts loosing its integrity(27). Therefore we decided test the system further by increasing the size ratio to 84% where LNPs with size 20*σ* have higher asphericity values.

#### Particle Size ratio: 84%

The highest particle-to-matrix size ratio examined in this study is approximately 84%, for LNP of size 20*σ* in the matrix mesh size of 24*σ*. As the LNP size increases, it faces significant barriers due to the matrix structure, but the total number of deformation modes also increases with the surface area. Figure 10 a) shows the evolution of mean square displacements of LNPs with the time lag. Here, the particle MSDs are always below 6000 *σ*^2^, suggesting fewer LNPs crossing the periodic boundaries of the simulation domain as compared to 50% and 67% size ratio cases. In this case, soft LNP shows the highest diffusive performance throughout the entire simulation. The MSD curve for the semi-elastic LNPs lies exactly in the midddle of the soft and hard LNP curves. The trend in the mean values for soft LNP remains linear till 1.7 × 10^5^ time lags which offers better reliability for our diffusivity predictions. Whereas for semi-elastic LNPs, the mean values shows little undulations about the linear trend within the fitting window. The curve for hard LNPs evolves perfectly in a straight line throughout the simulation span. Figure 10 b) presents the decaying nature of particle diffusivities in the matrix as their rigidity increases. Here, the soft LNPs (5.36 × 10^−3^*σ*^2^/*τ*) show almost threefold higher diffusivity compared to hard LNPs (1.83 × 10^−3^*σ*^2^/*τ*). Whereas the diffusivity of semi-elastic LNPs (3.32 × 10^−3^*σ*^2^/*τ*) is nearly twice that of the hard LNPs. This monotonic trend indicates the deformation-assisted diffusive transport of LNPs in the matrix environment.

**Figure 10.**
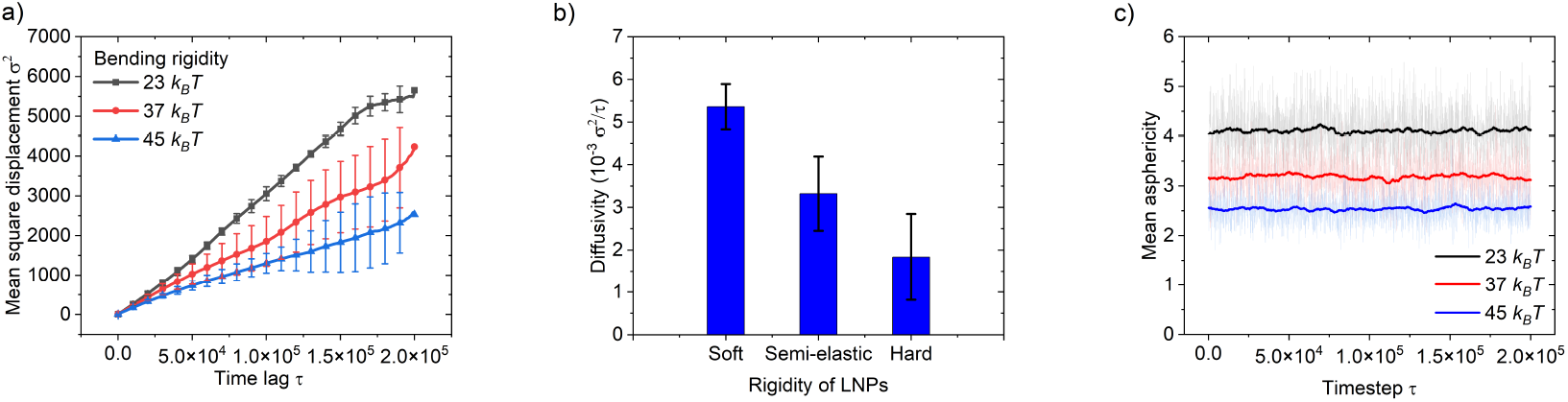
a) Mean square displacements, b) Diffusivity of LNPs with variable rigidities, and c) Mean asphericities of all LNP types for the size ratio of 84%.

Similarly, we compute the mean asphericities of the LNPs to visualize the deformation-induced shape change as they traverse in the matrix. Figure 10 c) shows the highest asphericity of 4 for the soft LNPs (23*k*_*B*_*T*), indicating maximum deviation from the spherical shape. Whereas, the semi-elastic (37*k*_*B*_*T*) and hard LNPs (45*k*_*B*_*T*) have mean asphericities around 3.2 and 2.5, respectively. For the higher size ratio of 84%, the degree of asphericity is strongly correlated with the diffusivity of the LNPs through the matrix. These observations confirm that the deformability of LNPs is influential in dictating diffusivity at a higher particle to matrix size ratios as compared to lower size ratios conditions. Both the particle size and the bending rigidity collectively influence the asphericity of particles within the matrix, which subsequently impacts the diffusivity of particles.

Lipid nanoparticle-based drug delivery is a promising platform for therapeutic molecules that are prone to degradation in the body. Despite of this, its efficacy is limited due to the multitude of effects that impact the diffusion of these nanoparticles in the biological microenvironment, such as the brain ECM. Softer particles often exhibit prolonged circulation times in the biological structures but they perform poorly in cellular internalization in the target cells(35). This common observation is convoluted by the concurrent changes in the effective particle to matrix size ratio, surface properties, and target site properties such as tumors cells. Particle deformability is one of the important properties in the design of nanoparticle based therapeutics, but its effect is difficult to quantify in isolation(24). These effects must be interpreted within the combined landscape of structural heterogeneity, surface chemistry, and size that collectively govern the multiscale biotransport of nanoparticles. The computational framework in this study offers a tool for studying the complex landscape of nanoparticle physicochemical properties and their impact on nanomedicine performance. Our model shows a way to disentangle the effect of the size ratio from deformation-assisted diffusion transport. The particle rigidity influences diffusive transport processes at higher size ratios as compared to the lower size ratio conditions. These insights inform the researchers to optimize the formulation process parameters that can reduce or increase the lipid nanoparticle size based on a particular composition and typical size ratios at the target site. The present framework can be extended for a more comprehensive evaluation of the effect of surface chemistry, heterogeneous mesh sizes, and target-dependent matrix stiffness. Such computational frameworks facilitate ranking between the key parameters of nanoparticles that dictate their performance in various regimes of their properties.

## CONCLUSION

In this study, we have demonstrated the collective role of lipid nanoparticle size and bending rigidity in the diffusive transport through the matrix. Our coarse-grained molecular dynamics simulations reveal that at lower size ratios such as 50% and 67%, the rigidity of particles does not influence their diffusive performance within the matrix. However, at higher size ratio of 84%, the diffusivity of particles decays with the rise in the bending rigidity. This result highlights the enhanced performance of soft LNPs as the particle-to-matrix mesh size ratio increases. These findings also underscore the critical role of particle size ratio in governing the diffusive transport of LNPs. Although this study is limited to matrices with uniform mesh size, in realistic biological environments, LNPs traverse through spatially heterogeneous matrices where varying mesh sizes dynamically alter their effective size ratio. In addition to this, these results address the role of the particle shape asphericity parameter that modulates the diffusive transport of LNPs in the matrix. This parameter increases with the particle size and decays with the bending rigidity of the particle. Therefore, they have a collective impact on the diffusivity of particles in the matrix. Our coarse-grained in-silico model for the LNP and the extracellular matrix system provides a robust framework to systematically study the impact of various physicochemical properties of LNPs on their diffusive performance. We note that in addition to particle size and rigidity, chemical properties such as particle–matrix affinity and surface charge also influence LNP performance, and will be investigated in the future. The findings of our study help in disentangling the effect of particle size and rigidity on the diffusive transport in the matrix. Along with this, it effectively complements the rational experimental design of the lipid nanoparticle-based drug delivery platforms involving complex biological matrices present in the brain parenchyma.

## AUTHOR CONTRIBUTIONS

P.N. and A.M.A. designed the research. P.N. carried out simulations, analyzed the data, and wrote the original draft. A.M.A. reviewed and edited the manuscript, analyzed the data, supervised the research, and acquired the funding.

## ACKNOWLEDGMENTS

We gratefully acknowledge the financial support from the Eli Lilly and Company.

