## Supplementary Materials for "Modeling lipid nanoparticle transport in extracellular matrix: Effects of particle size and rigidity"

### Domain dependence test:

To test the effect of system size on the diffusive behavior of LNPs in the matrix, we tested cases with size 98, 122, 146, 170, and 194$\sigma$. Table S1 lists the details of these cases. Here, all the model parameters are kept constant.

Table S1: Details of the cases performed for domain dependence test

| Domain size | Number of crosslinking nodal beads in each direction | Number of LNPs | Total bead count (LNPs & matrix) |
|---|---|---|---|
| 98 | $4 \times 4 \times 4$ | 10 | 7224 |
| 122 | $5 \times 5 \times 5$ | 20 | 14325 |
| 146 | $6 \times 6 \times 6$ | 34 | 24452 |
| 170 | $7 \times 7 \times 7$ | 54 | 38809 |
| 192 | $8 \times 8 \times 8$ | 80 | 57600 |

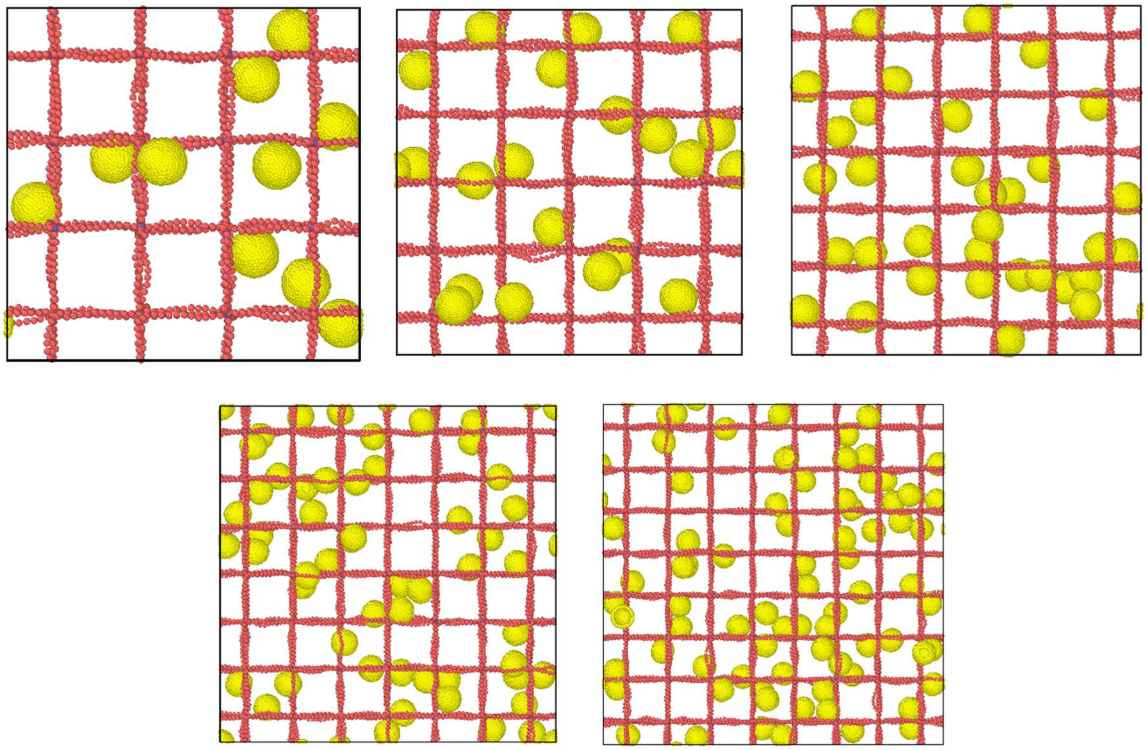

Figure S1: Snapshots of the five cases performed for domain dependence test

### Bending rigidity parameter $\mu$:

To tune the bending rigidity of LNPs in our system, we refer to the parametric curve presented in Figure S2 reproduced from data in Ref. [1]. We then choose three representative values of μ for different lipid nanoparticle types soft, semi-elastic, and hard.

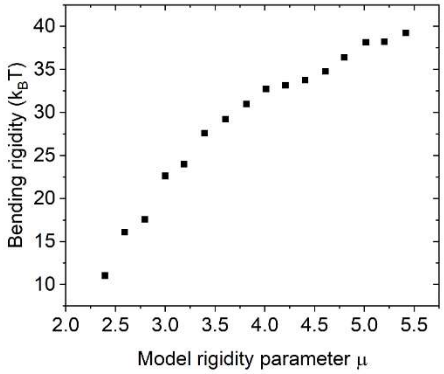


Figure S2: Change in the membrane bending rigidity with the parameter μ reproduced from data in Ref. [1]

### Fitting MSD curve for diffusivity:

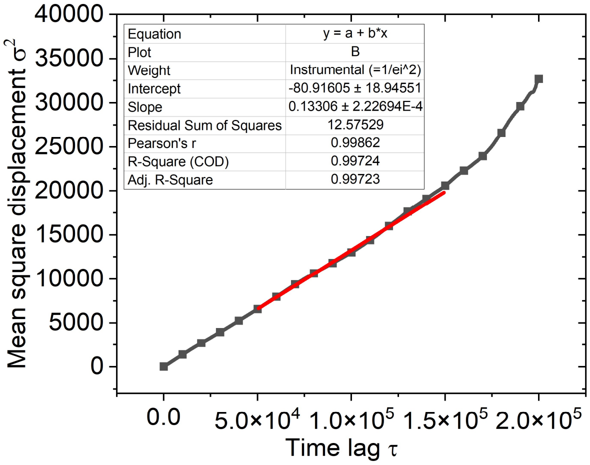


Figure S3: Linear fit of the MSD vs time lag using the period between 50,000 $\tau$ to 150,000 $\tau$

For fitting the MSD, we use the portion of MSD between time lags 50,000 $\tau$ to 150,000 $\tau$ as shown in the Figure S3. At shorter time lags, the motion of particles is predominantly due to the thermal fluctuations/vibrations. This region of the MSD curve doesn't capture the translational motion of particles in the matrix. Therefore, we neglect this part in our diffusivity calculations. On the other hand, at longer time lags the displacements of particles are of translational nature. But in this region, the mean square displacements are averaged over smaller samples that are separated by longer time lags. Due to this, there is an increased uncertainty in the MSD curve at longer time lags. Here, we avoid both of these regions and choose the middle 50% portion of the MSD curve for the diffusivity calculation ($k/6$), where $k$ is the slop of the linear fit.

#### Asphericity calculation:

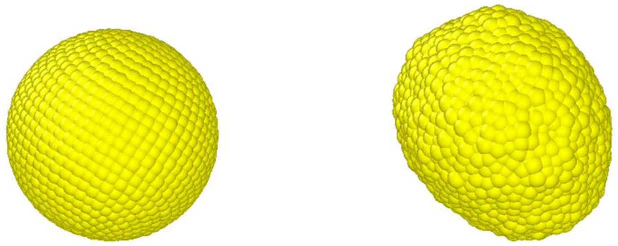

Figure S4: Deviation of soft LNP shape from perfect sphere to an elongated structure.

To quantify the deviation of LNP shape we use asphericity parameter $b$. We calculate this parameter from the eigenvalues of gyration tensor generated from the coordinates of the beads that form the LNP structure. Here, $b = \lambda_1 - \frac{1}{2}(\lambda_2 + \lambda_3)$ , $\lambda_1 > \lambda_2 > \lambda_3$ are the eigenvalues of gyration tensor.

In Figure S4, we show the perfectly spherical LNP on the left that gets deformed into an elongated shape on the right. In this example, we have soft LNPs (23$k_B$T) with size of 20$\sigma$ used in the case of 84% size ratio. For the perfect spherical shape all eigen values are equal to $33.333$, that makes asphericity parameter $b = 0$. For the elongated LNP shape on the right, $\lambda_1 = 36.012$, $\lambda_2 = 30.375$, and $\lambda_3 = 25.958$ which results in asphericity $b = 7.846$. This calculation is performed for all the LNPs in the system at each recorded time frame. We first take the ensemble average of the asphericity values over all the LNP population at individual timeframe and then take a moving time average of the values over 100 frames for the entire simulation runs. This approach provides an ensemble behavior of the LNP shapes over the duration of the simulation runs. In Figure S5, we show the evolution of asphericity values for particles in the case of size ratio 84%. For soft LNPs (23$k_B$T), the moving time averaged value is around $b = 4$, for the entire simulation span. The faded lines in the background show the ensemble average asphericity of LNPs at individual times over 2000 frames.

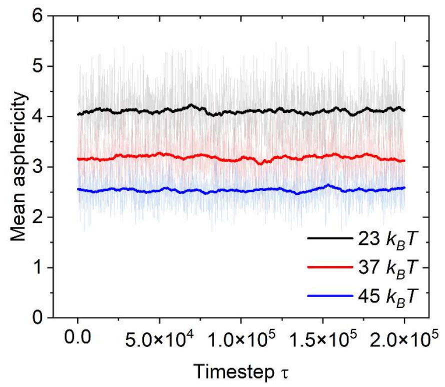


Figure S5: Time averaged mean asphericity of all LNP types used in size ratio 84%.

#### Relative diffusivity calculation:

We define the relative diffusivity as the ratio of the diffusivity of nanoparticles in the matrix to the diffusivity in water. Here, choose the hard LNPs to compare model performance against the experimental data. To calculate the diffusivity of hard LNPs in water, we performed cases without any matrix elements for different LNP sizes. Figure S6 shows the MSD curves obtained for LNPs in water for sizes 12, 16, and 20$\sigma$.

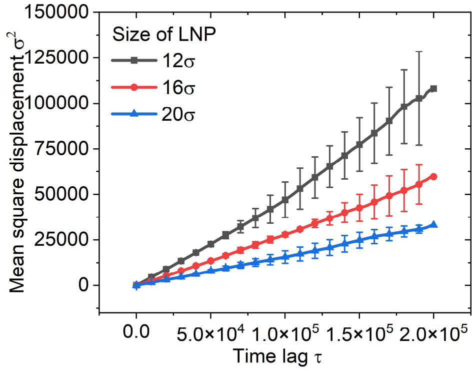


Figure S6: MSD curves for hard LNPs in water for different sizes.

We use the diffusivity data from hard LNP in the matrix and normalize against the respective diffusivities in water as shown below

Table S2: Relative diffusivity calculation for hard LNPs.

| Size ratio | Diffusivity in the matrix $(10^{-3}\sigma^2/\tau)$ | Diffusivity in water $(10^{-3}\sigma^2/\tau)$ | Relative diffusivity |
|---|---|---|---|
| 50% | $23.5 \pm 1.33$ | 85.98 | $0.273 \pm 0.015$ |
| 67% | $8.167 \pm 1.3$ | 48.33 | $0.169 \pm 0.027$ |
| 84% | $1.832 \pm 1.01$ | 27.10 | $0.068 \pm 0.037$ |
